# An integrative RNA spliceosomic landscape of pancreatic neuroendocrine tumors unveils novel clinicomolecular associations

**DOI:** 10.1101/2024.10.30.621040

**Authors:** Ricardo Blázquez-Encinas, Víctor García-Vioque, Andrea Mafficini, Luca Landoni, María Trinidad Moreno-Montilla, Nicolas Alcala, Sebastián Ventura, Eduardo Eyras, Salvatore Paiella, Roberto Salvia, Vita Rovite, Matthieu Foll, Claudio Luchini, Lynnette Fernandez-Cuesta, Rita T. Lawlor, Aldo Scarpa, Alejandro Ibáñez-Costa, Sergio Pedraza-Arevalo, Justo P. Castaño

**Affiliations:** Maimonides Institute for Biomedical Research of Cordoba (IMIBIC), Córdoba, Spain; Department of Cell Biology, Physiology, and Immunology, University of Córdoba, Córdoba, Spain; Reina Sofia University Hospital, Córdoba, Spain; Department of Engineering for Innovation Medicine, Section of Innovation Biomedicine, University and Hospital Trust of Verona, Verona, Italy; ARC-Net Research Centre, University and Hospital Trust of Verona, Verona, Italy; Department of General and Pancreatic Surgery, The Pancreas Institute, University and Hospital Trust of Verona, Verona, Italy; Rare Cancers Genomics Team (RCG), Genomic Epidemiology Branch (GEM), International Agency for Research on Cancer/World Health Organization (IARC/WHO), Lyon 69008, France; Department of Computer Sciences, University of Córdoba, Córdoba, Spain; The John Curtin School of Medical Research, The Australian National University Canberra Australian Capital Territory Australia.; EMBL Australia Partner Laboratory Network at the Australian National University Canberra Australian Capital Territory Australia.; Latvian Biomedical Research and Study Centre, Riga, Latvia; Department of Diagnostics and Public Health, Section of Pathology, University and Hospital Trust of Verona, Verona, Italy; CIBER Fisiopatología de la Obesidad y Nutrición (CIBERobn), Córdoba, Spain

**Keywords:** Pancreatic Neuroendocrine Tumors, PanNET, RNA splicing

## Abstract

Alterations in alternative splicing are emerging as a novel hallmark in cancer biology, offering new insights. However, integrative analyses of splicing are still scarce, particularly in rare cancers such as pancreatic neuroendocrine tumors (PanNETs). These tumors are highly heterogeneous, complicating diagnosis and treatment. This study is the first to comprehensively investigate the RNA splicing landscape in PanNETs, identifying distinct spliceosomic profiles correlated with unique clinical and molecular characteristics. We analyzed RNA-seq data from 174 samples, identifying three distinct spliceosomic groups (SPN1, SPN2, SPN3) with unique clinical and molecular characteristics. SPN1 exhibited intermediate clinical features and specific splicing machinery profile, SPN2 was associated with frequent mutations in *MEN1* and *DAXX*/*ATRX* genes, and SPN3 showed a prevalence of well-differentiated tumors with distinct splicing patterns. These groups were linked to different clinical outcomes and activated key biological processes like mTOR signaling and hormone secretion pathways. Our findings underscore the significant impact of RNA splicing on PanNET heterogeneity and suggest that detailed splicing profiles could serve as valuable tools for identifying novel biomarkers and therapeutic targets. This study provides crucial insights into PanNET molecular biology and paves the way for personalized therapies based on splicing features.

## INTRODUCTION

It is now widely accepted that alterations in alternative splicing relevantly contribute to cancer by impacting it at various levels, from tumor initiation to progression, metastasis, and treatment resistance [1–4]. Evidence to support this notion initially derived from reports showing the oncogenic role of individual altered components of the splicing machinery, like mutated or over/under-expressed splicing factors, or abnormal splice variants (reviewed in [1–3, 5]). Further, global *omics* analyses documented alterations in splicing machinery components, splicing events and/or isoforms in the main types of cancer [6–10]. Yet, to fully understand how altered splicing actually contributes to the features and behavior of a given cancer, it will be necessary to acquire a precise, quantitative picture of the status of the splicing machinery, splice events and transcript variants in that particular cancer, i.e., its *spliceosomic landscape*. This will enable examining the relationships among its underlying molecular mechanism and identify relevant clinicomolecular associations, with the ultimate goal of translating this information into precision medicine. While this could be more feasible in common cancers, it becomes particularly difficult for rare tumors such as pancreatic neuroendocrine tumors (PanNETs), a peculiar neoplastic category originating from the endocrine cells of the pancreas [11–13].

PanNENs display unique biological features and a clinical behavior that range from indolent and slow-growing to highly aggressive metastatic tumors, which coupled to their rising incidence [14] and remarkable heterogeneity [15] makes their diagnosis and clinical management a significant challenge [11, 12, 16–18]. The WHO classification categorizes PanNENs into well-differentiated neuroendocrine tumors (PanNETs), further divided into G1, G2 and G3, based on their mitotic count and Ki-67 index, and poorly differentiated G3 carcinomas (PanNECs) [11, 19, 20]. Depending on their hormone-secreting activity, PanNETs are classified into functional and non-functional tumors [11]. Their only curative approach, when feasible, is surgery, while current pharmacological treatments include somatostatin analogs, mTOR pathway inhibitors, DNA repair pathways inhibitors and antiangiogenic drugs [11–13, 17, 18].

Collaborative studies have provided comprehensive molecular landscapes of PanNETs, including genomic [21], transcriptomic [22, 23], proteomic [24], and DNA-methylomic [25] profiles. Most commonly mutated genes in PanNETs are *MEN1* and *DAXX/ATRX*, with less frequent mutations in PI3K/mTOR pathway genes, such as *TSC2*, *PIK3CA*, and *PTEN* [21]. As in other neoplasms, transcriptomic profiling has led to identify molecular signatures that correlate with tumor grade, prognosis, and therapeutic response in PanNETs [22–24]. In particular, categorizing them based on their mRNA and miRNA expression profiles uncovered distinct subtypes such as the insulinoma-like (IT) and metastasis-like primary (MLP) PanNETs [22, 26]. However, despite these advances, the molecular mechanisms underlying PanNETs behavior remain poorly understood and translation of this molecular information to the clinical practice is very limited [16, 27, 28]. In this scenario, recent evidence that alterations in RNA biology and alternative splicing represent a novel and informative cancer hallmark has opened new research avenues to better understand PanNETs [29]. Indeed, in recent studies we discovered that the splicing machinery is severely dysregulated in PanNETs as compared to non-tumor tissue, identifying two splicing factors, *NOVA1* [30] and *CELF4* [31], that are overexpressed in tumors, and associated to oncogenic features and treatment response. Similar findings have been reported in different cancers, including pancreatic adenocarcinoma, where alterations of splicing factors have been associated to metastasis or cancer development [1–4, 32]. However, to date, a comprehensive analysis of RNA splicing has not been conducted in PanNETs. Here, we have analyzed clinical, molecular and transcriptomic data from a large cohort of 174 tumors to undertake the first integrative spliceosomic landscape of PanNETs. This approach revealed the existence of separate PanNET spliceosomic groups displaying distinct splicing-related clinic-molecular associations, providing novel information to achieve a more nuanced understanding of the intricate, multilayered underpinnings of splicing alterations in these rare heterogeneous tumors.

## MATERIAL AND METHODS

### Sample collection

The cohort included 174 well-differentiated pancreatic neuroendocrine tumor (PanNET) cases. A set of 20 non-neoplastic samples were also retrieved and used as a reference for variant calling. Patient samples and data were collected by the ARC-Net Research Centre biobank, University of Verona, Italy, under Prog. 1885 (Prot. N. 52070) from the Integrated University Hospital Trust Ethics Committee (AUOI-CE) and used in this study under protocol CE2172 (Prot. N. 26773). All samples were collected from surgical resections, snap frozen and stored at -80°C before further processing. Diagnosis and related clinicopathological data were verified by two experienced pancreatic pathologists (AS, CL). The data produced during the current study is available from the corresponding author upon reasonable request.

### DNA and RNA extraction

For both DNA and RNA extraction, tumor samples had full face frozen sectioning performed in optimal cutting temperature (OCT) medium. DNA was isolated using the QiAamp DNA Mini Kit (Qiagen, Milan, Italy) according to the manufacturer instructions. RNA was isolated using the miRNeasy Mini Kit (Qiagen, Milan, Italy) according to the manufacturer instructions. Assessment of both DNA and RNA was performed using a Qubit and Bioanalyzer to confirm quantitation and quality, respectively, as previously reported [33].

### DNA sequencing and analysis

Targeted DNA sequencing was performed using the SureSelectXT HS CD Glasgow Cancer Core assay (Agilent, Santa Clara, CA), hereinafter referred to as CORE. The assay was developed in collaboration with the Glasgow Precision Oncology Laboratory (https://www.gpol.org) based on a comprehensive analysis of data from previously performed whole-genome sequencing [34]. The panel targets ∼1.8 megabases of the genome to investigate somatic mutations, copy number alterations and structural rearrangement in 174 genes with proven involvement in cancer development and therapy. Sequencing libraries were prepared by targeted capture from 70 ng of genomic DNA using the SureSelect HT kit and the SureSelect Enzymatic Fragmentation Kit (Agilent, Santa Clara, CA) following the manufacturer’s instructions. Quality and quantity of purified pre-capture libraries was assessed using the Qubit BR dsDNA assay (ThermoFisher, Milan, Italy). Hybridization-capture and purification of the libraries was performed according to the manufacturer’s instructions, using 1.6 µg of pooled DNA from 16 pre-capture libraries. 100 ng from each pre-capture library was used to prepare the 16-library pools. Captured library pools were enriched by PCR, purified and quantified using the Qubit dsDNA HS assay. Quality of the library pools was verified with the Agilent 4200 Tape Station and High Sensitivity D1000 ScreenTape (Agilent, Santa Clara, CA). Sequencing was performed on a NextSeq 500 (Illumina, San Diego, CA) loaded with 2 captured library pools, using a high-output flow cell and 2×75bp paired end sequencing.

Paired FASTQ files were aligned to the human reference genome (version hg38/GRCh38) using BWA and saved in the BAM file format [35]. BAM files were sorted, subjected to PCR duplicate removal and indexed using biobambam2 v2.0.146 [36]. Coverage statistics were produced using samtools [37]. Variant calling was performed on tumor samples using a set of 20 non-neoplastic samples as a reference. Single nucleotide variants were called using Shearwater [38]. Small (<200 bp) insertions and deletions were called using Pindel [39]. Single nucleotide variants were further annotated using a custom pipeline based on vcflib (https://github.com/ekg/vcflib), SnpSift [40], the Variant Effect Predictor (VEP) software [41] and the NCBI RefSeq transcripts database (https://www.ncbi.nlm.nih.gov/refseq/). Annotated variants were filtered keeping only missense, nonsense, frameshift, or splice site variants as annotated from the canonical transcripts. In case of alternative exon usage between transcripts, the transcript where the variant produced the worst effect was retained and used for annotation. All candidate mutations were manually reviewed using Integrative Genomics Viewer (IGV), version 2.4 [42] to exclude sequencing artefacts.

### RNA sequencing

Libraries for RNAseq were prepared using 100ng of total RNA with the TruSeq Stranded Total RNA Sample Prep Kit, Ribo-Zero Gold rRNA Removal and unique dual indices according to the manufacturer’s instructions (Illumina, San Diego, CA). The concentration and size distribution of the libraries was assessed with the Agilent Bioanalyzer (Santa Clara, CA) and Qubit fluorimeter (Invitrogen, Carlsbad, CA). Sequencing was performed following Illumina’s standard protocol either using a) the Illumina cBot and HiSeq 3000/4000 PE Cluster Kit or b) the Illumina NovaSeq 6000 and an S2 flow cell. For the HiSeq, 100 × 2 paired end reads were sequenced on an Illumina HiSeq 4000 using Hiseq 3000/4000 sequencing kit and HCS v3.3.52 collection software. Base-calling was performed using Illumina’s RTA version 2.7.3. For the NovaSeq, 150 × 2 paired end reads were sequenced using the NovaSeq S2 reagent kit and NovaSeq Control Software v1.3.1. Base-calling was performed using Illumina’s RTA version 3.3.3.

### RNA-seq processing and alignment

The raw sequencing reads were first subjected to quality control using FastQC to assess data quality and identify potential issues. The reads were then aligned to the human reference genome (GRCh38) using the STAR aligner [43]. STAR was chosen for its superior performance in aligning RNA-seq reads and its ability to effectively handle splicing events. Alignment parameters were set to default, with adjustments made for specific experimental requirements as necessary.

### Quantification of gene expression

Following alignment, the generated BAM files were used to quantify gene expression levels. Python tool *htseq-count* [44] was employed to count the number of reads mapping to each gene based on the overlap with annotated genes in the GRCh38.

### Differential expression analysis

The raw counts were then normalized to account for differences in library size and sequencing depth among samples using DESeq2 [45]. Differential expression analysis was conducted to identify genes with statistically significant changes in expression across different groups of samples. Genes were considered differentially expressed based on a threshold of fold-change > 2 and an adjusted p-value < 0.05, correcting for multiple testing using the Benjamini-Hochberg method. To identify genes that define each of the groups, pair comparisons were performed, and the intersection of the genes overexpressed in a group compared to the other two were selected.

### RNA splicing analysis

rMATS-turbo [46] was employed to systematically investigate alternative splicing events across the samples. rMATS is a computational tool designed for the robust detection and quantification of splicing events from RNA-seq data by calculating Percent Spliced In (PSI). PSI quantifies the relative abundance of a specific splicing event in each sample. The tool analyzes RNA-seq reads to identify and quantify PSI of the five major types of splicing events: skipped exons (SE), retained introns (RI), mutually exclusive exons (MXE), alternative 5’ splice sites (A5SS) and alternative 3’ splice sites (A3SS). We used the –novelSS flag to identify novel splice sites and we set the significance threshold for differential alternative splicing events at FDR < 0.05 and |dPSI| > 0.1. The rMATS output, which includes detailed annotations of splicing events and their quantitative changes between conditions, was further analyzed.

### Unsupervised clustering

Raw reads from htseq-count were imported into R and normalized using variance stabilizing transformation (VST) function of DESeq2. VST counts were obtained from 315 splicing machinery related genes, selected by the method described by Paschalis *et al* [47]. Unsupervised clustering of the samples was made using the VST counts of the 315 splicing machinery genes, by Ward’s clustering method with Euclidean clustering distance, using *pheatmap* R package. Clustering with the different types of RNA splicing events (SE, RI, MXE, A5SS and A3SS) were performed using PSI from most variable events (those that explain 50% of the variance across the samples), which were: 3054 for SE, 386 for RI, 10662 for MXE, 490 for A5SS, and 632 for A3SS.

### Enrichment analysis

Gene enrichment analysis was made with differentially expressed genes and differentially spliced genes, using *clusterProfiler* R package [48], and obtaining Gene Ontology: Biological Process and KEGG databases from the Molecular Signatures Databases (MSigDB). Significative pathways were considered with *p* < 0.01. To calculate biological pathways scores, single-sample Gene Set Enrichment Analysis (ssGSEA) was made using *GSVA* R package [49].

### Tumor deconvolution

Infiltration of cells from tumor microenvironment was estimated from RNA-seq counts using *EPIC* R package [50]. This package estimates proportion of immune, stromal and endothelial cells from bulk RNA-seq data. Transcripts per million (TPM) were calculated to use the *EPIC* function, and proportion of Cancer Associated Fibroblasts (CAFs), macrophages, endothelial cells, CD4 and CD8 T cells and B cells was calculated for each of the samples.

### Splice events characterization

rMATS output files with splicing events quantification for each sample were further analyzed using two different tools: ExonOntology [51], that uses exons genomic coordinates to infer protein features; and SpliceTools [52], that analyze splice events characteristics. As ExonOntology only works with hg19 genome annotations, we converted them from rMATS files, using UCSC liftover (minimum ratio of bases that must remap: 0.95). Sashimi plots of specific alternative splicing events (ASE) were performed using rmats2sashimiplot visualization tool (https://github.com/Xinglab/rmats2sashimiplot).

### Partial Least Squares Discriminant Analysis (PLS-DA)

To dissect the complex interplay between splicing machinery gene expression and the delineation of spliceosomic groups, we employed PLS-DA utilizing the *mixOmics* package in R [53]. This multivariate statistical approach was chosen for its efficacy in handling high-dimensional data and its ability to maximize the variance between predefined groups while considering the correlation structure within the data. Following model construction, Variable Importance in Projection (VIP) scores were computed for each splicing machinery gene, identifying those with the most significant contribution to the discrimination between groups.

### Motif enrichment analysis

Motif enrichment was performed using the sequences of excluded SE events of each group, including 400 bp intronic regions adjacent to them, and comparing with the sequences of included SE events of that same group (including 400 intronic adjacent bp). These analysis were performed using Simple Enrichment Analysis (SEA) tool [54] from The MEME Suite [55].

### Statistical analysis

All statistical analyses performed were done in R.

## RESULTS

### The expression profile of the splicing machinery delineates three distinct groups of PanNETs

Spliceosomic focused analysis was started by employing the levels of expression of a predefined set of genes integral to the splicing machinery to generate an unsupervised clustering of 174 PanNETs samples. This approach unveiled three distinct groups that we termed Splicing PanNET1 (SPN1), SPN2 and SPN3, which displayed a clearly unique expression pattern of the splicing machinery and, most importantly, were associated with divergent clinical and molecular features (**Fig 1A**). Thus, while SPN1 and SPN2 comprised similar proportions of G1 and G2 PanNETs and lacked G3 tumors, SPN3 was marked by a predominance of grade 1 PanNETs (Fisher’s exact test, *p* = 8.27e-03; **Fig 1B**), and, most notably, exclusively contained all G3 PanNETs. Ki-67 proliferation index was significantly elevated in SPN3 PanNETs in comparison to SPN2 (**Fig 1C**), likely linked to the inclusion of G3 samples in SPN3. In fact, excluding G3 samples from the analysis revealed a significantly higher Ki-67 index in SPN2 PanNETs (**Supp Fig 1A**), and, at the gene expression level, *MKI67* exhibited higher expression in SPN2 samples even when including G3 (**Supp Fig 1B**). Moreover, an increased frequency of metastatic PanNETs patients was observed in SPN2 (Fisher’s exact test, *p* = 3.05e-02; **Fig 1D**). An examination of common PanNETs mutations revealed a significant concentration of *MEN1* mutations in SPN2 (Fisher’s exact test, *p* = 6.47e-09; **Fig 1E**), with the lowest frequency of mutations in SPN3, while SPN1 exhibited an intermediate frequency. Likewise, analysis of *DAXX* and *ATRX* mutations further identified a significant overrepresentation of mutated tumors in SPN2 for both genes (Fisher’s exact test, *p* = 5.17e-10 and *p* = 3.56e-03, respectively, **Fig 1E**). These results indicate that clustering PanNETs using a spliceosomic approach reveals three distinct groups of tumors displaying specific clinical and molecular features that may not be solely explained by their histological grade.

**Fig 1.**
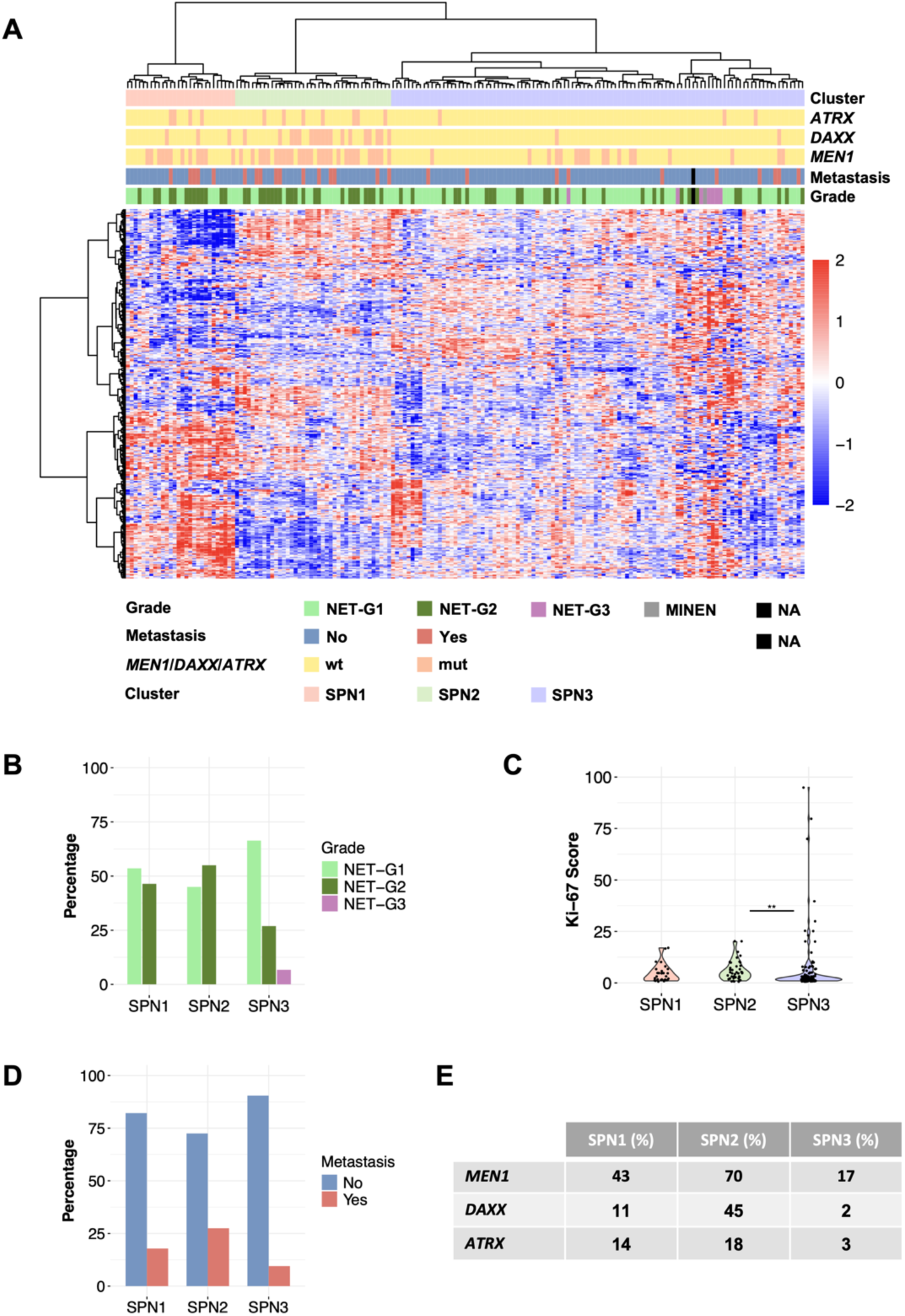
Unsupervised spliceosomic clustering of PanNETs reveals 3 groups with distinct clinico-molecular features. **A** Heatmap illustrates the unsupervised clustering of 174 PanNET based on the expression levels of the 315 genes of the splicing machinery across individual samples, with samples grouped according to the identified spliceosomic clusters. Presence of mutations in *MEN1*, *ATRX* and *DAXX*, metastasis and histological grade are indicated. **B** Frequency of distribution of PanNET grades across spliceosomic groups. Fisher’s exact test indicated enrichment of G1 in SPN3 samples (*p* = 8.27e-03). **C** Comparison of the Ki-67 proliferation index among the spliceosomic groups. Asterisks indicate statistical significance of Dunn’s tests (** *p* < 0.01). **D** Distribution of presence/absence of metastasis across spliceosomic groups. Fisher’s exact test indicated enrichment of metastatic samples in SPN2 (*p* = 3.05e-02). **E** Frequencies of mutations in *MEN1*, *ATRX* and *DAXX* genes across spliceosomic groups. Fisher’s exact test respectively indicated enrichment of mutated *MEN1*, *ATRX* and *DAXX* samples in SPN2 (*MEN1*: *p* = 6.47e-09, *ATRX*: *p* = 3.56e-03, *DAXX*: *p* = 5.17e-10).

### The spliceosomic groups have distinct transcriptomic signatures

To further delineate the molecular underpinnings of the three spliceosomic PanNET groups we conducted a differential gene expression analysis to explore their specific transcriptomic signatures. By employing pairwise comparisons and identifying genes uniquely overexpressed in each group relative to the others, we uncovered distinct molecular pathways characterizing each group (see **Supp Table 1**). Interestingly, KEGG pathway analysis revealed a substantial disparity in the types of pathways associated to each spliceosomic subtype. Thus, while SPN1 is marked uniquely by an upregulation of protein export and spliceosome-related genes, SPN2, in contrast, is distinguished by an enriched metabolism of cell membrane lipids, like sphingolipid and arachidonic acid, and immune-related pathways, like neutrophil extracellular trap formation or cytokine-cytokine receptor interaction. On the other hand, SPN3 shows a number of selectively enriched pathways, with a particular preponderance of heightened signaling related to hormone production, secretion, and synapse-related processes (**Fig 2A**). The basis for some of these findings may be related with the absence of functional tumors in SPN2, as opposed to the presence of 18.5% and 11.7% functional tumors in SPN1 and SPN3, respectively (**Fig 2B**). Nevertheless, whether some of the differences observed among SPN groups are associated with or even functionally linked to their distinct spliceosomic profiles awaits elucidation.

**Fig 2.**
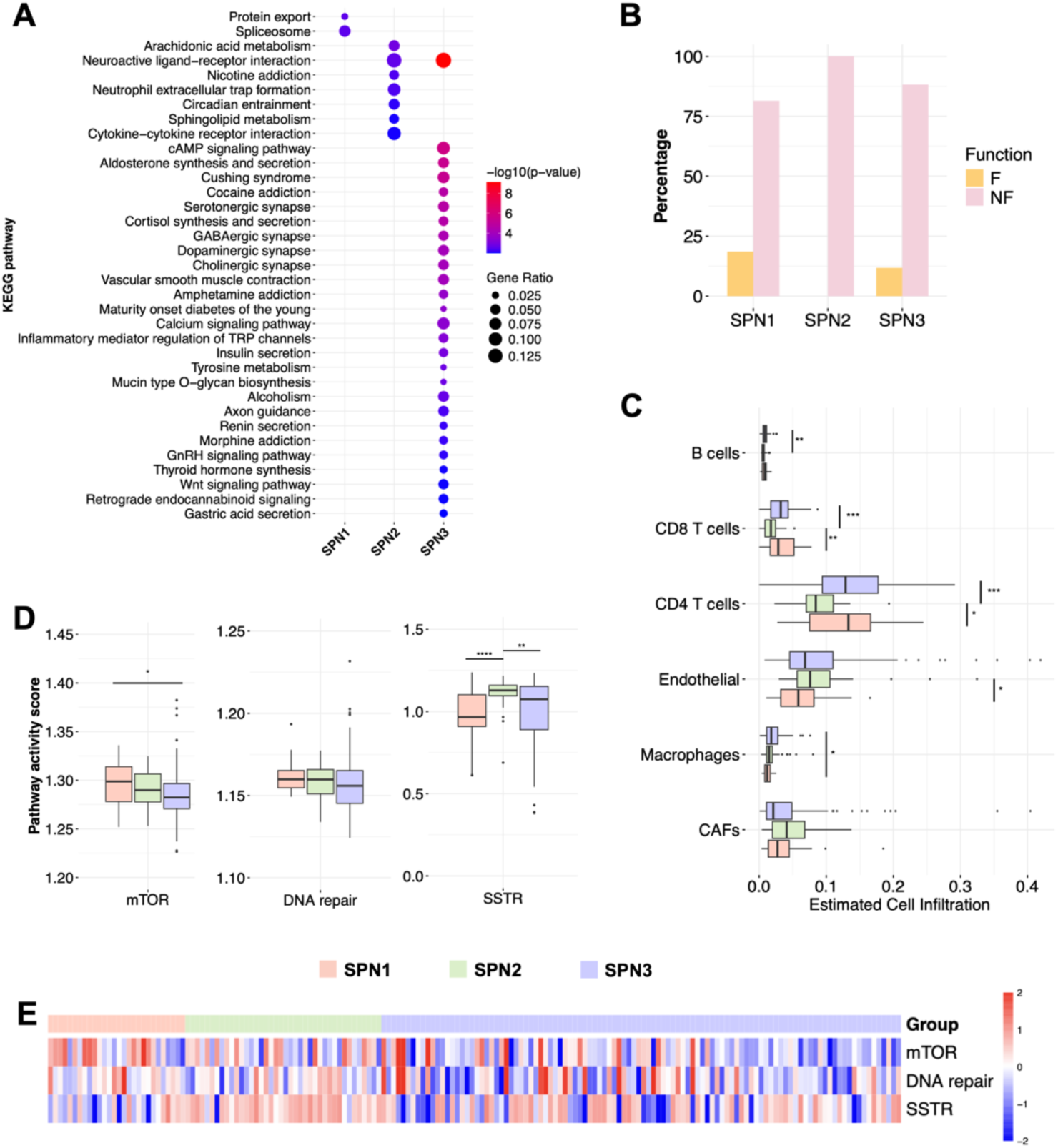
**A** KEGG pathway analysis highlighting distinct biological pathways enriched in each spliceosomic group. The size of the spheres indicates gene ratio, which is ratio of genes affected by total genes in that pathway. The color of the spheres indicates the logarithm of the *p* value. **B** Distribution of functional/nonfunctional samples across spliceosomic groups. Notice that SPN2 only includes nonfunctional samples. **C** Box plot illustrating the estimated proportions of various tumor microenvironment cells, including B cells, CD4 and CD8 T cells, endothelial cells, macrophages and CAFs across the spliceosomic groups, as inferred by the EPIC algorithm. **D** Box plots showing the ssGSEA scores for mTOR, DNA repair and SSTR pathways activation across the three spliceosomic groups. Asterisks in C-D indicate statistical significance of Dunn’s tests (* *p* < 0.05, ** *p* < 0.01, *** *p* < 0.001, **** *p* < 0.0001). **E** Heatmap showing heterogeneity in scores of mTOR, DNA repair and SSTR pathways activation across the samples of each spliceosomic group.

To delve deeper into the complex nature of the spliceosomic groups, we explored whether they differed in the proportion of microenvironment immune cells, as estimated by EPIC (**Fig 2C**). This revealed that SPN1 exhibits reduced levels of macrophages and endothelial cells, while SPN2 shows lower levels of CD4 and CD8 T lymphocytes and B lymphocytes. While the underlying causes of this differential immune cell infiltration among spliceosomic groups is not known, it unveils a varying tumor microenvironment across the groups that may entail relevant implications.

To explore the potential links between splicing and key clinically related molecular targets in PanNETs, we assessed the activation of relevant biologically targetable pathways by calculating single-sample GSEA scores for the SSTR, mTOR, and DNA repair pathways (**Fig 2D**). SPN1 displayed the highest activation of the mTOR pathway, significantly surpassing SPN3. DNA repair pathways did not show significant differences among the groups. Notably, SSTR pathway activation was markedly higher in SPN2 compared to the others, although this activation varied within SPN1 and SPN3 samples, indicating heterogeneity within these groups (**Fig 2E**). These findings reveal a previously unrecognized, pathway-selective association between the expression profile of the splicing machinery and clinically relevant targets.

### The pattern of alternative splicing events was distinct for each spliceosomic group

Having examined the molecular engine responsible for alternative splicing, we then focused on the analysis of its resulting outcome, i.e., the splice variants, by quantitatively comparing the distinctive patterns of ASEs across the groups. First, by categorizing and quantifying the Percent Spliced In (PSI) for each of the 5 main ASE types (**Fig 3A**) —skipped exons (SE), retained introns (RI), mutually exclusive exons (MXE), and alternative 5’ and 3’ splice sites (A5SS and A3SS)—we observed that SE was the most prevalent ASE, followed by MXE, A3SS, A5SS, and RI (**Fig 3B**).

**Fig 3.**
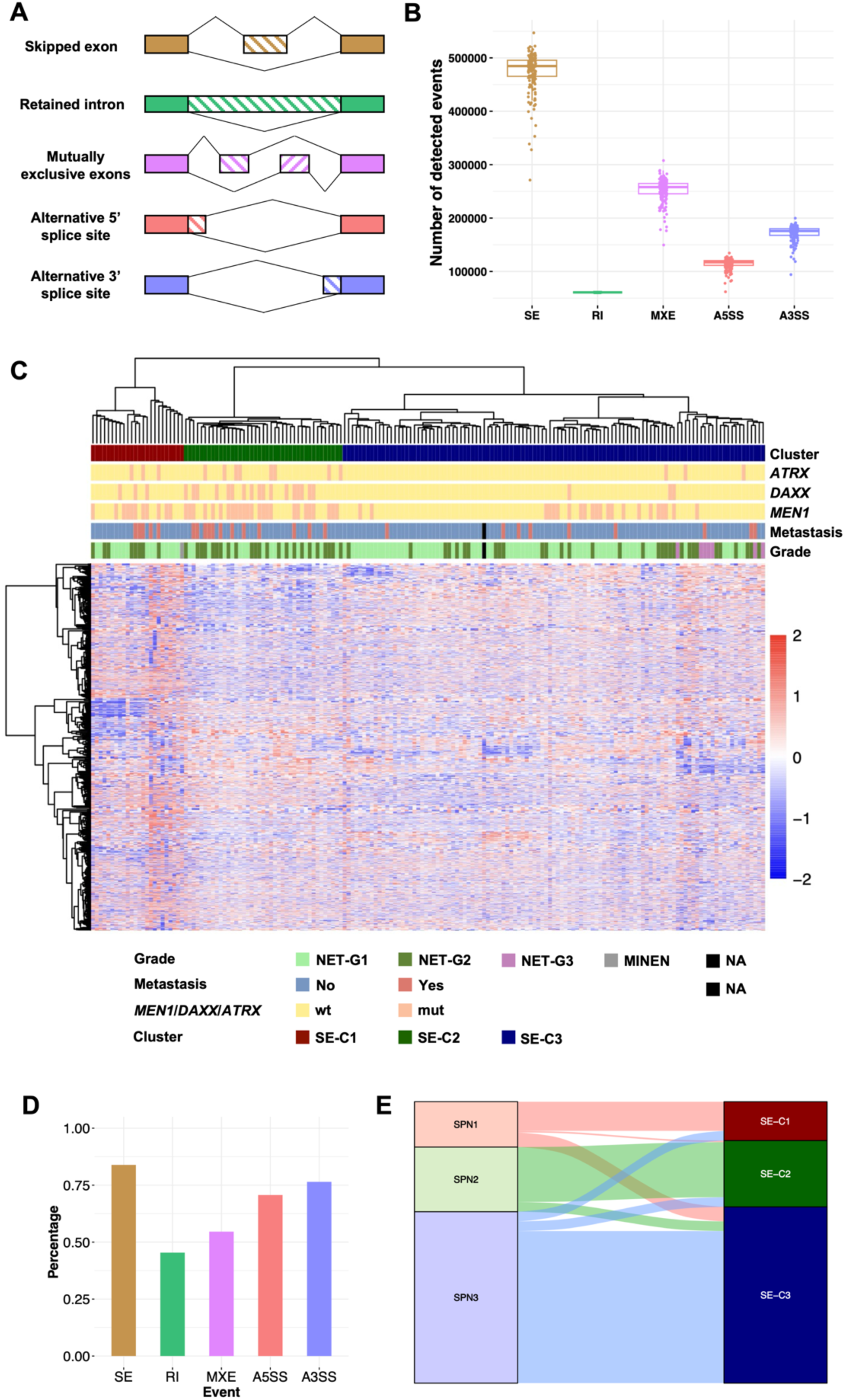
**A** Graphical representation of the five major types of alternative splicing events (ASEs) analyzed: Skipped Exons (SE), Retained Introns (RI), Mutually Exclusive Exons (MXE), Alternative 5’ Splice Sites (A5SS), and Alternative 3’ Splice Sites (A3SS). **B** Box plot depicting the frequency of each ASE type detected in the PanNET samples. **C** Heatmap showing the results of unsupervised clustering of PanNET samples using PSI from the most variable SE ASEs across the samples. Grade, presence of metastasis and mutations in *MEN1*, *ATRX* and *DAXX* are indicated. **D** Bar graph illustrating the proportion of samples that consistently group together in the clustering analyses based on splicing machinery expression profiles and the most variable ASEs by type of event. Fisher’s exact test respectively indicated association of splicing machinery and the distinct splice events clusterings (SE: *p* < 2.20e-16, RI: *p* = 2.28e-09, MXE: *p* = 5.94e-14, A5SS: *p* < 2.20e-16, A3SS: *p* < 2.20e-16). **E** Alluvial plot showing the correlation between the clustering outcomes based on splicing machinery expression and the most variable SE ASEs.

Then, we mirrored the unsupervised clustering approach followed with the expression of the splicing machinery genes but applying it, instead, to the most variable ASE across the PanNETs samples, i.e., for each ASE type, we selected those that explained 50 % of the variance across the samples (**Fig 3C, Supp Fig 2**). As depicted by the heatmaps, this approach resulted, using any of the five types of ASE, in the generation of three major clusters, as in the case of the splicing machinery, (e.g. those derived from SE events in **Fig 3C**, named SE-C1, SE-C2 and SE-C3). Actually, a closer examination of their clinicomolecular features hinted that these clusters display visible similarities and overlaps with the spliceosomic groups discovered earlier (SPN1, SPN2 and SPN3; **Fig 1A**). Indeed, the level of alignment of samples in the same group between the two clustering methods were: 83.9% for SE, 45.4% for RI, 54.6% for MXE, 70.7% for A5SS and 76.4% for A3SS (**Fig 3D**). This congruence was further validated through Fisher’s exact test, statistically substantiating the high, visually evident correspondence between the SPN groups and those generated by analyzing ASE, illustrated in the alluvial plots, which confirmed that particularly SE (**Fig 3E**), but also A5SS and A3SS clustering, had the highest equivalence with the SPN spliceosomic groups (SE: *p* < 2.20e-16, RI: *p* = 2.28e-09, MXE: *p* = 5.94e-14, A5SS: *p* < 2.20e-16, A3SS: *p* < 2.20e-16; **Supp Fig 2**). These results offer primary evidence to support the notion that the distinctive patterns of ASE in PanNETs share a remarkable level of coincidence with the transcriptomic profiles determined by the levels of expression of the components of the splicing machinery (i.e., the SPN groups).

### Molecular deepening to characterize differential ASE among groups

The unique correspondences between the SPN and ASE spliceosomic clusters prompted us to conduct an in-depth analysis to identify differential ASEs between groups (**Supp Table 2-4**). Interestingly, differential ASE followed specific patterns depending on the comparison made, as, if we compared SPN1 vs SPN2 or vs SPN3, the most common differential ASE type was MXE. However, when comparing SPN2 vs SPN3, SE was the most common type of differential ASE (**Supp Fig 3**). Further, by intersecting ASEs that were significantly differential in both pairwise comparisons involving each group, we delineated a unique ASE profile for each group. For example, for SPN1, we used those events that were significantly differential against SPN2 and also against SPN3. In this sense, the profile of ASE was distinct for each of the spliceosomic groups (**Fig 4A**), with MXE events being more frequent in SPN1, and SE events in SPN2 and SPN3. Mean PSI was calculated for all exons and introns that could be skipped or retained, showing that general inclusion of exons was higher in SPN3 than in SPN2. Also, SPN1 had the highest inclusion of introns, followed by SPN3, being SPN2 the one with lowest intron retention (**Fig 4B**). Of note, these ASE that were intrinsic to each group affected different biological pathways. In SPN1, ASE affected genes mainly involved in ribosome, autophagy, protein degradation, sphingolipid and redox pathways (**Fig 4C**). Meanwhile, in SPN2, ASE affected genes that belonged to cell signaling (specially insulin mediated) and lipid metabolism pathways (**Fig 4D**). In SPN3, ASE affected the splicing machinery itself, and also ribosome and motor proteins related genes (**Fig 4E**). We visualized the alteration of some ASE affecting relevant genes using Sashimi plots (**Fig 4F**), as the histone acetyltransferase component (*ELP2*), the centromere protein X (*CENPX*) or the nucleobindin 2 (*NUCB2*).

**Fig 4.**
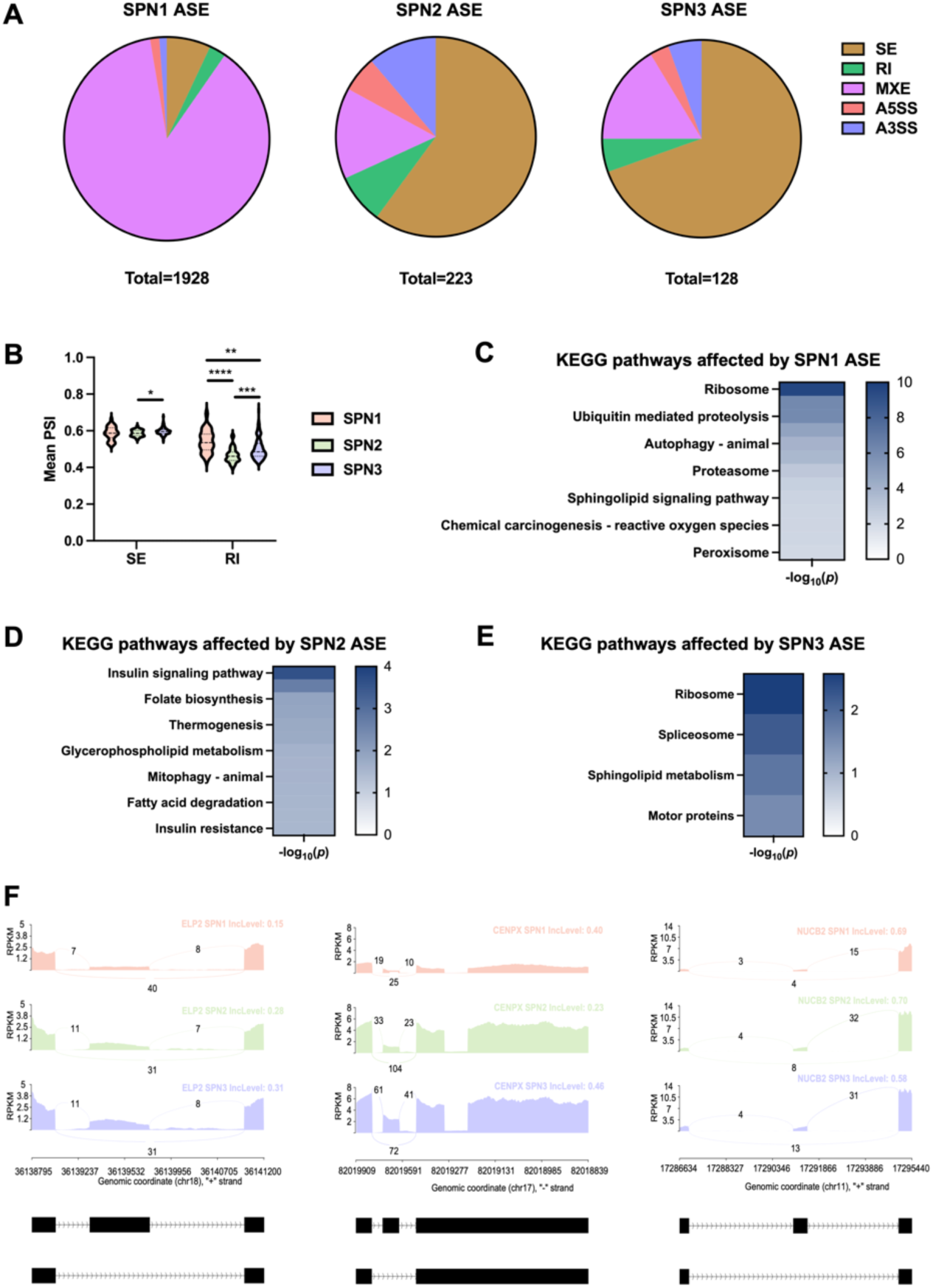
**A** Pie charts showing distribution of ASE types in the intrinsic ASE of each spliceosomic group (SPN1 at the left, SPN2 at the center and SPN3 at the right). **B** Comparison of the mean PSI of all the differential SE and RI events in the three spliceosomic groups. **C-E** KEGG pathways enrichment of genes affected by intrinsic ASE of SPN1 (**C**), SPN2 (**D**) and SPN3 (**E**). **F** Representative Sashimi plot showing SE events from each of the spliceosomic groups in relevant genes. The number of junctions reads and the Percent Spliced In (IncLevel) are indicated in the plot for SPN1 (orange), SPN2 (green) and SPN3 (blue).

We then analyzed SE differential events between pairwise comparisons in depth using two exploratory software: ExonOntology and SpliceTools. This revealed that differential SE events in SPN1 vs SPN2 comparison were similar to those in SPN1 vs SPN3. Interestingly, around 25 % of the SE included in SPN1 were not annotated previously (**Supp Fig 4A**). Also, more than one third of the events occurred outside the coding sequence (CDS; **Supp Fig 4B**). Most of these exons coded for post-translational modification (PTM) sites, being around a quarter of them phosphorylation sites (**Supp Fig 4C-D**). However, when comparing differential SE events between SPN2 vs SPN3, almost three-quarters of the exons were in the CDS (**Supp Fig 4B**), with similar annotation frequencies in each group (**Supp Fig 4A**). In this case, the most common annotation for these exons were structure domains for the protein (**Supp Fig 4C**), and about half of the exons affected phosphorylation sites (**Supp Fig 4D**). These findings further unveil unexpected characteristics associated to the spliceosomic features of PanNETs, suggesting, for example the existence of novel splicing isoforms, ASE that may not solely reside in the coding regions and ASE that could affect functionally relevant domains of the genes, all of which can be tumor-subtype specific.

An additional, relevant aspect examined was the sizes of the differential SE, which were, all of them, larger than the average of annotated exons in the genome (**Supp Fig 5A**). Interestingly, exons included in SPN2 were larger than in SPN1 and SPN3. However, the sizes of the upstream exons and introns of the SE events were larger for SPN1-included SE. Comparing splice sites scores unveiled that those exons included in SPN2 and SPN3 had lower scores in donor and acceptor splice sites that those exons included in SPN1 (**Supp Fig 5B**). Also, all the differential SE events had lower scores that the mean of annotated exons.

Some of the exon inclusions may lead the particular splice variants generated to be degraded by the nonsense-mediated decay (NMD). We estimated which exons were targeted for NMD among all differential SE events. More than a third of the exons included in SPN1 compared to the other groups were estimated to lead to NMD degradation (**Supp Fig 5C**). SPN3 has also higher proportion of NMD degradation than SPN2.

On the other hand, in the case of RI events, the introns involved in the differential events had all reduced sizes compared with annotated introns (**Supp Fig 5D**). Also, introns retained in SPN2 were the shortest, and introns retained in SPN1 the longest. Differential RI events had lower splice scores in donor and acceptor sites when comparing with the mean of annotated RI events (**Supp Fig 5E**).

### The splicing machinery expression profile is different for each spliceosomic group

Partial Least Squares Discriminant Analysis (PLSDA) was used to maximize the variance among the 3 groups generated by using the splicing machinery, SPN1, SPN2 and SPN3, and to determine which splicing machinery components were the most discriminant for each of the groups, showing separation of the 3 groups (**Fig 5A**). To determine which splicing machinery genes were mostly contributing to each of the groups, PLS-DA was performed with each group against the rest (**Supp Fig 6**), and the component associated to the group of interest was selected using t-tests (**Supp Fig 6**). Then, Variable Importance in Prediction (VIP) scores were calculated to understand the contribution of each gene belonging to the splicing machinery to each group according to the PLS-DA model. In this sense, top 10 splicing machinery genes which discriminate each group against the others were obtained (**Fig 5B, C, D**), unveiling genes as *LSM5*, *CACTIN* or *SMNDC1* for SPN1; *DDX1*, *PRPF38A* or *SF3B1* for SPN2; and *SNRPB2*, *SF3A1* or *SF3B4* for SPN3.

**Fig 5.**
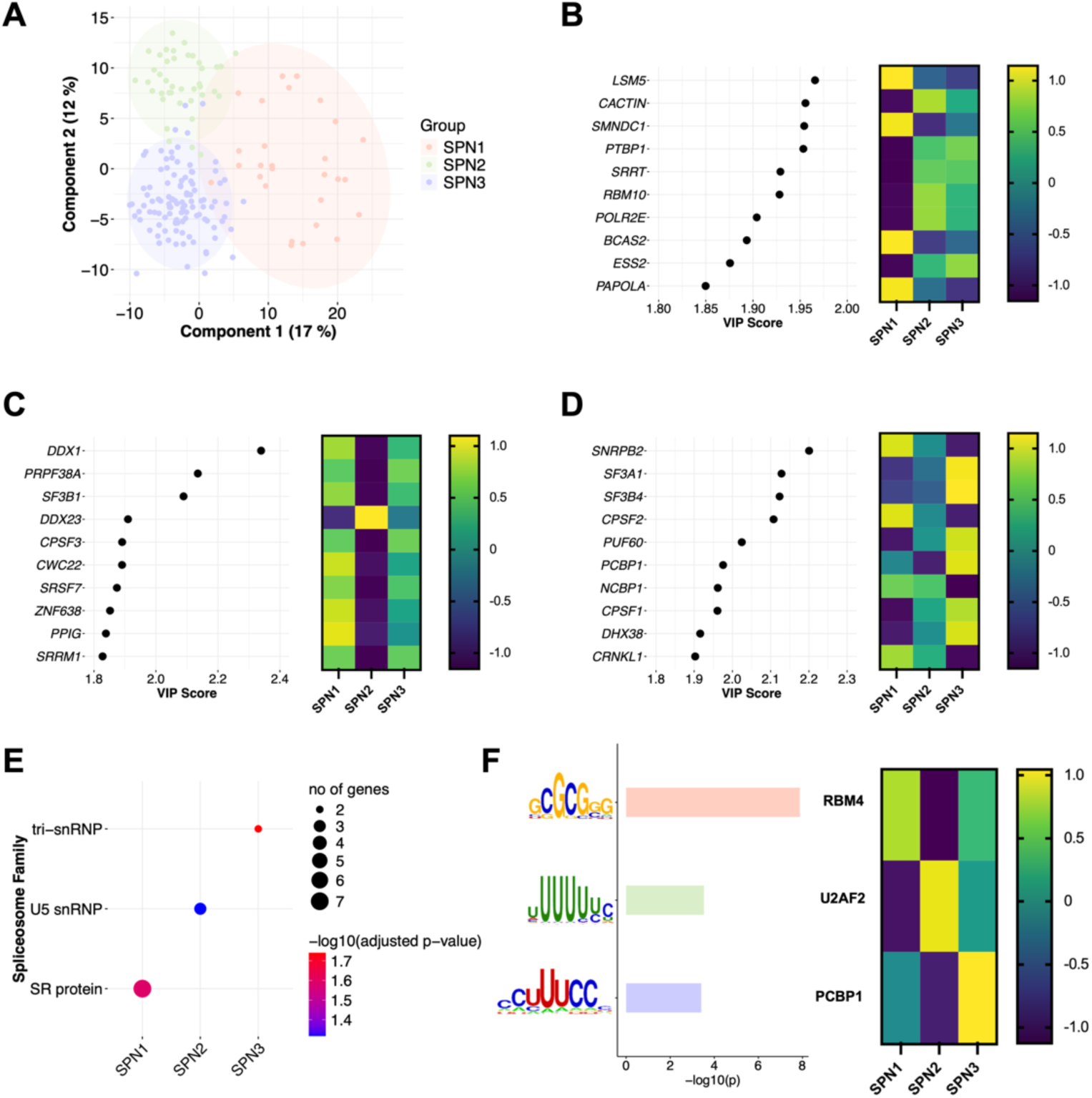
**A** Partial Least Squares Discriminant Analysis (PLS-DA) clustering showing the separation among the three spliceosomic groups based on the expression profiles of splicing machinery genes. **B-C-D** Dot plots presenting the top 10 splicing machinery genes with the highest Variable Importance in Prediction (VIP) scores for SPN1 (**B**), SPN2 (**C**), and SPN3 (**D**), respectively. Next to them, corresponding heatmaps showing mean expression of these splicing machinery genes in each of the groups. **E** Dot plot depicting the enrichment of specific splicing machinery functions in each spliceosomic group. The size of the spheres indicates number of genes of that family that were significant. The color of the spheres indicates the logarithm of the *p* value. **F** Enrichment of motifs found significant in each spliceosomic group (SPN1-orange, SPN2-green, SPN3-blue), when comparing excluded SE events vs included SE events in each group. The bar plot shows the logarithm of the *p* value. The heatmap show the mean expression of the predicted splicing machinery gene that joins that motif in each group.

The different components of the splicing machinery accomplish distinct functions in the RNA splicing process. To determine if there was an overrepresentation of a splicing machinery function in each spliceosomic group, we obtained those splicing machinery components that statistically correlated to the groups of PLSDA components (Pearson correlation test, *r* > 0.5, *p* < 0.05). We then used Fisher’s exact test to look for enrichment functions of the splicing machinery components of each group. This strategy revealed that SPN2 and SPN3 had core spliceosome components overexpressed (**Fig 5E**). In particular, SPN2 was characterized by U5 snRNP components overexpression and SPN3 had tri-snRNP components overexpressed. In contrast, SPN1 was characterized by overexpression of SR proteins, which are auxiliary factors in RNA splicing (**Fig 5E**), showing that each spliceosomic group was defined by different families within the splicing machinery.

Next, we applied motif enrichment analysis to SE events that were skipped in each group, compared to those that were included in the same group, obtaining a list of motifs for each of the groups, with their respective prediction of RNA-binding protein that recognizes that sequence (**Supp Table 5-7**). We looked for those splicing machinery components that were predicted to recognize each group motifs and that were overexpressed in that spliceosomic group. There was one significative motif in each spliceosomic group that fulfilled both criteria (GCGCGGG for SPN1, UUUUUUC for SPN2 and CCUUUCC for SPN3). These motifs were recognized by RBM4, U2AF2 and PCBP1, respectively; and these three splicing machinery components were overexpressed in their respective group, when compared to the others (**Fig 5F**). Collectively, these results suggest that the spliceosomic groups defined by the expression profile of the splicing machinery convey unique splicing-driven features that may entail relevant functional implications.

## DISCUSSION

Unlike other cellular functions whose machineries comprise a relatively lower and more enclosed set of components (e.g., translation by the ribosome), a considerable number of molecular players integratively interact with the spliceosome to complete the splicing process [56]. Accordingly, to generate a comprehensive and inclusive spliceosomic picture of this molecular engine, we decided to use the expression levels of 315 genes involved in the splicing machinery [47] to generate an unsupervised clustering. This strategy segregated PanNETs into three unique groups —SPN1, SPN2 and SPN3— that displayed distinct associations with molecular and clinical features of the tumors. Indeed, while it was not necessarily anticipated that the mutational profile of the spliceosomic groups differed, we found that SPN2 tumors displayed higher mutation frequencies in *MEN1*, *ATRX* and *DAXX*, whereas SPN1 showed intermediate frequencies. These observations can be relevant because *MEN1* and *ATRX*/*DAXX* losses are adverse prognostic factors in PanNETs [57, 58]. The role of ATRX/DAXX is related to alternative lengthening of telomeres [59], and their potential interaction with splicing alterations has not been examined in detail yet. However, such a link may exist, as we recently reported that *NOVA1*, a key splicing factor for pancreatic beta cells [60], is overexpressed and potentially oncogenic in PanNETs, where its silencing reduced expression of ATRX and DAXX in NET cell models [30]. Furthermore, menin, the product of the most frequently mutated gene in PanNETs, *MEN1*, has recently been described as a pivotal regulator of functionally relevant RNA splicing programs in lung cancer models, through mechanisms involving suppression of RNA polymerase II elongation and R-loop-induced genome instability [61]. Thus, in light of the previous evidence, our findings strongly suggest a potential functional association between these gene mutations and splicing dysregulation in PanNETs, whose putative pathobiological implications warrant further investigation.

Given the essential role of alternative splicing to dynamically model the transcriptional pattern of eukaryotic cells, both in health and disease [62, 63], it was conceivable that PanNET spliceosomic groups could display distinct transcriptomic profiles. Indeed, exploring their linked functional pathways revealed insightful associations. Thus, while the higher activation in protein export/secretion pathways in SPN1 and SPN3 may relate to the higher proportion of functional PanNETs, which have increased hormone secretion, whether there is an association between tumor functionality and spliceosomic profiles is not known and requires additional research. In this same line, we found clear differences in the relative abundance of the various cell populations comprising the tumor microenvironment among the spliceosomic groups, with lower proportion of T and B lymphocytes in SPN2, and higher proportion of endothelial cells. The relevance of the precise cell populations that integrate tumor microenvironment is undisputable, particularly in the context of immunotherapy [64]. The variability in immune cell infiltration across spliceosomic groups suggests that alternative splicing may influence, and/or be reciprocally modulated by, the tumor microenvironment. This original notion would open an interesting scenario, as it is actionable and may affect immune surveillance and response to therapy [65], or could even bring new opportunities to use immune-based treatments, which are being linked to splicing dysregulation [66]. The putative links of the spliceosomic groups may involve other relevant therapeutic targets in PanNETs, where SPN1 displayed increased signaling activation in mTOR and SPN2 in SSTR. The relationship between these PanNET clinical targets and splicing alteration was not totally unexpected, as we originally proposed that a disrupted splicing process could lie behind the seminal discovery of an oncogenic, aberrantly truncated variant of SST_5_, SST_5_TMD4 [67–73], which, in fact, prompted the subsequent demonstration that the machinery of splicing is dysregulated in PanNETs [30] and a number of tumors [29, 32, 74–78]. In turn, this led us to uncover a tight relationship between two of the most altered splicing factors in PanNETs, NOVA1 [30] and CELF4 [31], the regulation of the mTOR pathway, and the consequent response to drugs that, like everolimus, are currently used to treat NETs, although, to date, without reliable biomarkers to predict response [79]. These results shed novel light into the precise molecular mechanisms underlying splicing dysregulation in PanNETs, its functional implications and its relationship with the intrinsic heterogeneity of these tumors, which definitely impacts their variable response to treatments. A more profound comprehension of this complex interplay could leverage new strategies based on splicing profiles to improve tumor classification and prediction of disease progression and treatment response, and to move the goal of personalized medicine closer to PanNET patients.

The biological meaning and potential translational relevance of the complex associations among spliceosomic and clinicomolecular parameters identified here are not all readily evident and will require additional work. However, certain discoveries, like the striking overlap between groups generated through unsupervised clustering of splicing machinery expression and those derived from the most variable ASEs, can already offer relevant, original information. A recurrent difficult question in splicing research is what are the real and precise consequences that the changes in the machinery impart into splicing variant selection and its subsequent functional outcomes. While multiple studies have documented how individual defects in splicing machinery components (e.g., mutation in splicing factors) directly drive functionally relevant changes in specific splice variants, the precise influence of general splicing machinery patterns into functionally relevant alterations of splicing variants is still poorly defined. Indeed, even in PanNETs [30, 31], recent studies have associated the frequency of certain splice events to specific splicing machinery components. However, the fact that tumors with different profiles of splicing machinery expression have different ASE patterns had not been described before in PanNETs or, at least in this form, in other cancer types. Intriguingly, our results indicate that the association of ASE and splicing machinery groups is especially strong for SE. While the exact mechanisms underlying this observation remain unknown, it is noteworthy that the three groups present differences in the sizes and the sequences of the exons and introns that they differentially splice. Elucidating the underpinnings of these interrelationships can be relevant, in that alternative splicing changes affect essential protein features, including their structure and post-translational modifications sites, especially phosphorylation, crucial in signal transduction [80–83]. Moreover, alternative splicing might also affect the stability of RNA molecules, directing them to be degraded by the NMD, and thereby determining their translation into proteins [84, 85]. The functional implications of all these ASEs, could help explain the distinct biological pathways enriched in the different groups linked to each of the ASE, which, as observed for the SPN groups, involve pathways as important as protein metabolism, cell signaling, and the splicing machinery itself. Future studies are warranted to explore in detail the functional relevance of the multiple splice variants identified in this study, many previously unknown. Evidence for their potential significance is provided by three representative examples of splice variants previously reported to be involved in pancreatic cancer risk (*ELP2* [86]), MYC cancer dysregulation (*CENPX* [87]) or insulin secretion/glucose metabolism (*NUCB2* [88]).

Another observation worth considering is the association of spliceosomic groups with specific families of splicing machinery components. Previous studies revealed a clear dysregulation of the splicing machinery in PanNETs and identified specific components that could provide novel therapeutic targets [30, 31]. Here, we show that families of splicing machinery components’ may distinguish PanNET spliceosomic groups, potentially serving as biomarkers for their functional differentiation. Inasmuch as these families subserve disparate roles in splicing, their association with each spliceosomic group, i.e., the auxiliary SR protein family with SPN1, and the core components in SPN2 and SPN3, will likely render different outcomes in the ASE and, hence, the resulting splice variants. In line with this idea, we discovered specific sequences that were enriched in motifs that are targets for specific splicing machinery components (e.g., RBM4, U2AF2 and PCBP1) that belong to their respective, expected families.

This study has some limitations, given the challenge posed by the complexity of RNA splicing mechanisms and their regulation, which preclude to readily elucidate the functional consequences of many identified ASEs. Thus, experimental studies will be needed to delineate the functional impacts of specific ASEs and splicing factors identified in this study, particularly their roles in tumor growth, progression, and response to therapy. As well, replicating this approach in a comparable set of PanNETs, albeit recognizably difficult, would provide further support and wider insights into our current findings.

In conclusion, the comprehensive RNA splicing analysis presented here constitutes the first approach to acquire a true spliceosomic landscape of PanNETs. Our results unveil the existence of distinct, previously unrecognized spliceosomic groups displaying markedly distinct clinicomolecular features, which can provide a novel framework to better understand the molecular heterogeneity of PanNETs, offering new avenues for research and clinical management of this complex disease. Furthermore, the striking correspondence discovered between spliceosomic groups defined by the splicing machinery and those generated on the basis of specific splicing events uncover an intriguing link with novel scientific and pathobiological implications for these rare tumors, which should also be explored in other cancers.

## FUNDINGS

This work was supported by Spanish Ministry of Economy [MINECO; M (JPC)] and Ministry of Science and Innovation [MICINN; PID2019-105201RB-I00, AEI/10.13039/501100011033 (JPC)]. Spanish Ministry of Universities Predoctoral contracts FPU18/02275 (RBE) and FPU20/03958 (VGV). Junta de Andalucía (BIO-0139); FEDER UCO-202099901918904 (JPC and AIC). Postdoctoral contract under the program María Zambrano funded by the European Union Next Generation-EU (SPA). Asociación Cáncer de Páncreas (ACAPAN) and AESPANC 2022 (JPC and AIC). Grupo Español de Tumores Neuroendocrinos y Endocrinos (GETNE2016 and GETNE2019 Research grants; JPC). Fundación Eugenio Rodríguez Pascual (FERP2020 Grant; JPC). CIBERobn Fisiopatología de la Obesidad y Nutrición. CIBER is an initiative of Instituto de Salud Carlos III. Part of this work was supported by COST (European Cooperation in Science and Technology – www.cost.eu) through the COST Action TRANSPAN (CA21116). Associazione Italiana Ricerca sul Cancro (AIRC IG number: 26343); Fondazione Cariverona: Oncology Biobank Project ‘Antonio Schiavi’ (prot. 203885/2017); Fondazione Italiana Malattie Pancreas (FIMP-J38D19000690001); European Union - NextGenerationEU through the Italian Ministry of University and Research under PNRR - M4C2-I1.3 Project PE_00000019 “HEAL ITALIA” to AS CUP: B33C22001030006.

## CONFLICT OF INTEREST

Where authors are identified as personnel of the International Agency for Research on Cancer/WHO, the authors alone are responsible for the views expressed in this article and they do not necessarily represent the decisions, policy, or views of the International Agency for Research on Cancer/WHO.

## DATA AVAILABILITY

The raw data underlying this article are available in European Genome-Phenome Archive at https://ega-archive.org, and can be accessed with accession number EGAS50000000670.

---

**Supp Fig 1.**
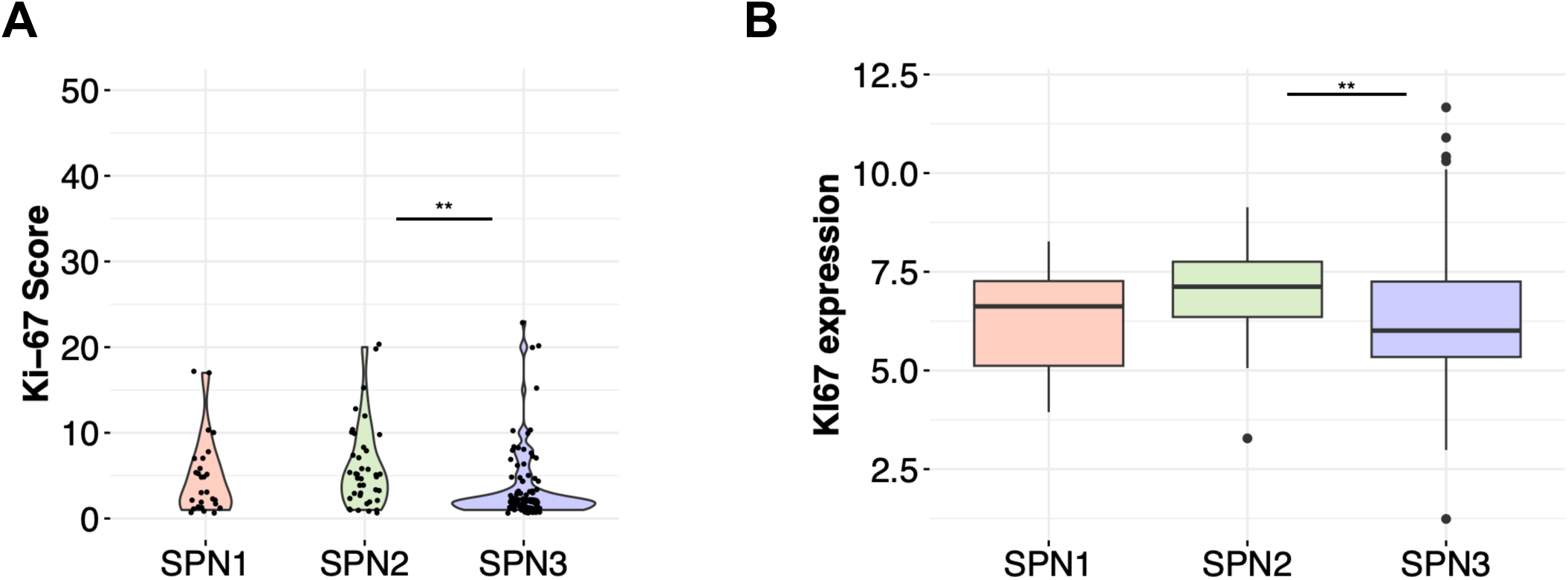
**A** Comparison of the Ki-67 proliferation index among the 3 spliceosomic groups, SPN1, SPN2 and SPN3, without including grade 3 (G3) samples. **B** Comparison of *MKI67* expression levels among the 3 spliceosomic groups SPN1, SPN2 and SPN3 (including G3). Asterisks indicate statistical significance of Dunn’s tests (** *p* < 0.01).

**Supp Fig 2.**
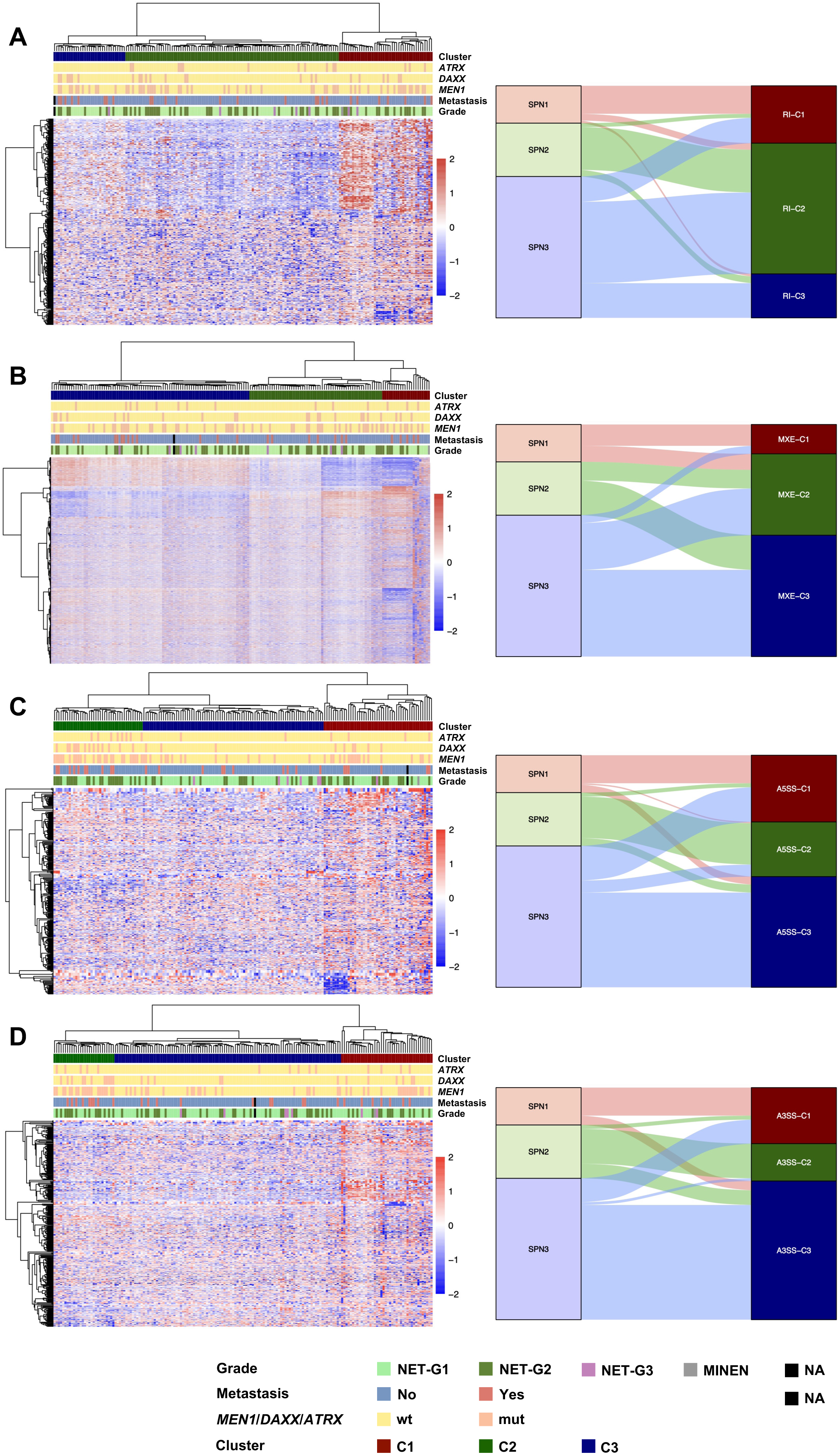
Heatmaps showing the results of unsupervised clustering of PanNET samples using PSI from the most variable ASEs across the samples (left). Alluvial plot showing the correlation between the clustering outcomes based on splicing machinery expression and the most variable ASEs (right). **A** represents clustering with RI events, **B** for MXE events, **C** for A5SS events and **D** for A3SS events.

**Supp Fig 3.**
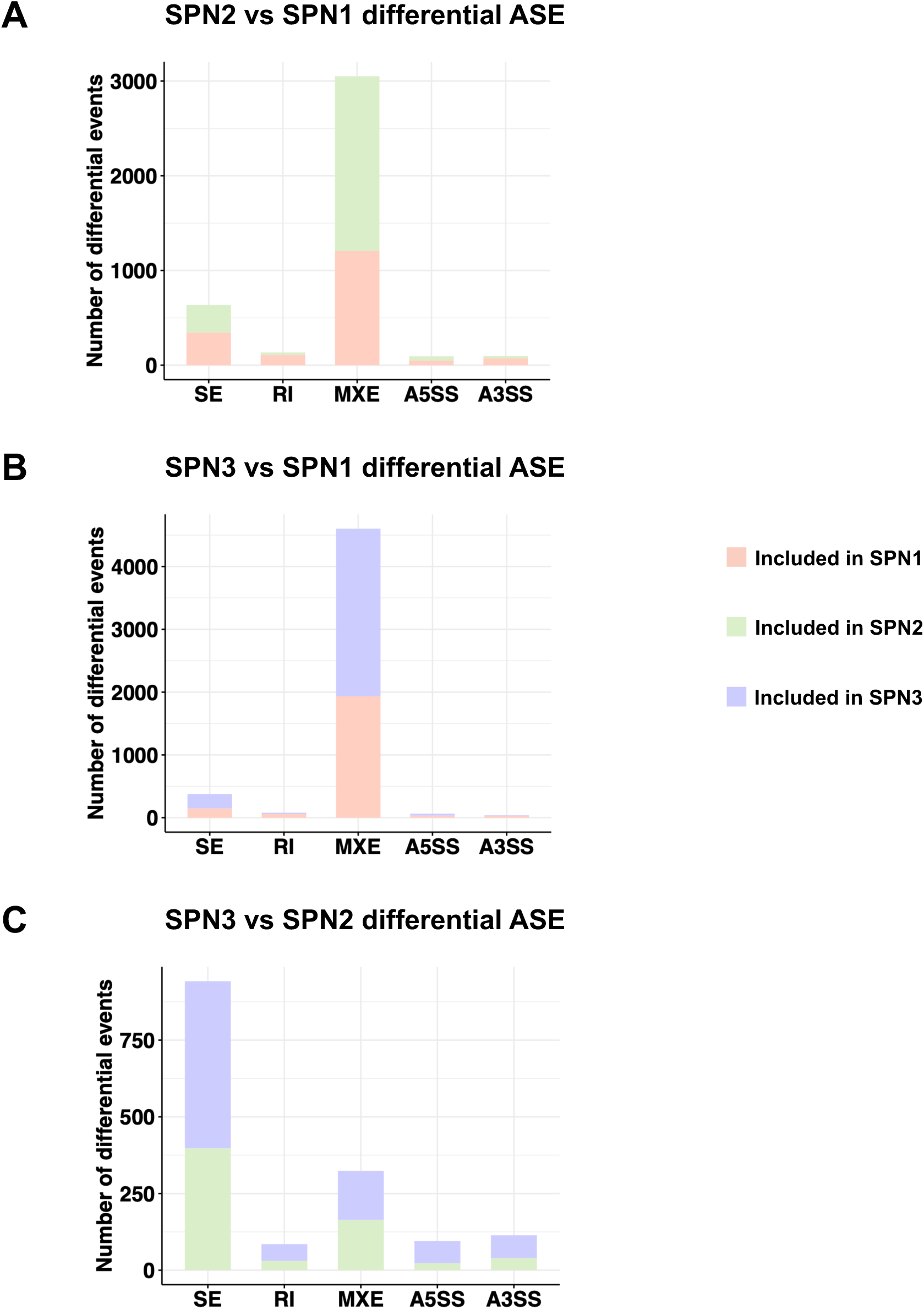
Bar plot showing number of differential alternative splicing events by type (SE, RI, MXE, A5SS and A3SS) when comparing SPN2 vs SPN1 (**A**), SPN3 vs SPN1 (**B**), and SPN3 vs SPN2 (**C**).

**Supp Fig 4.**
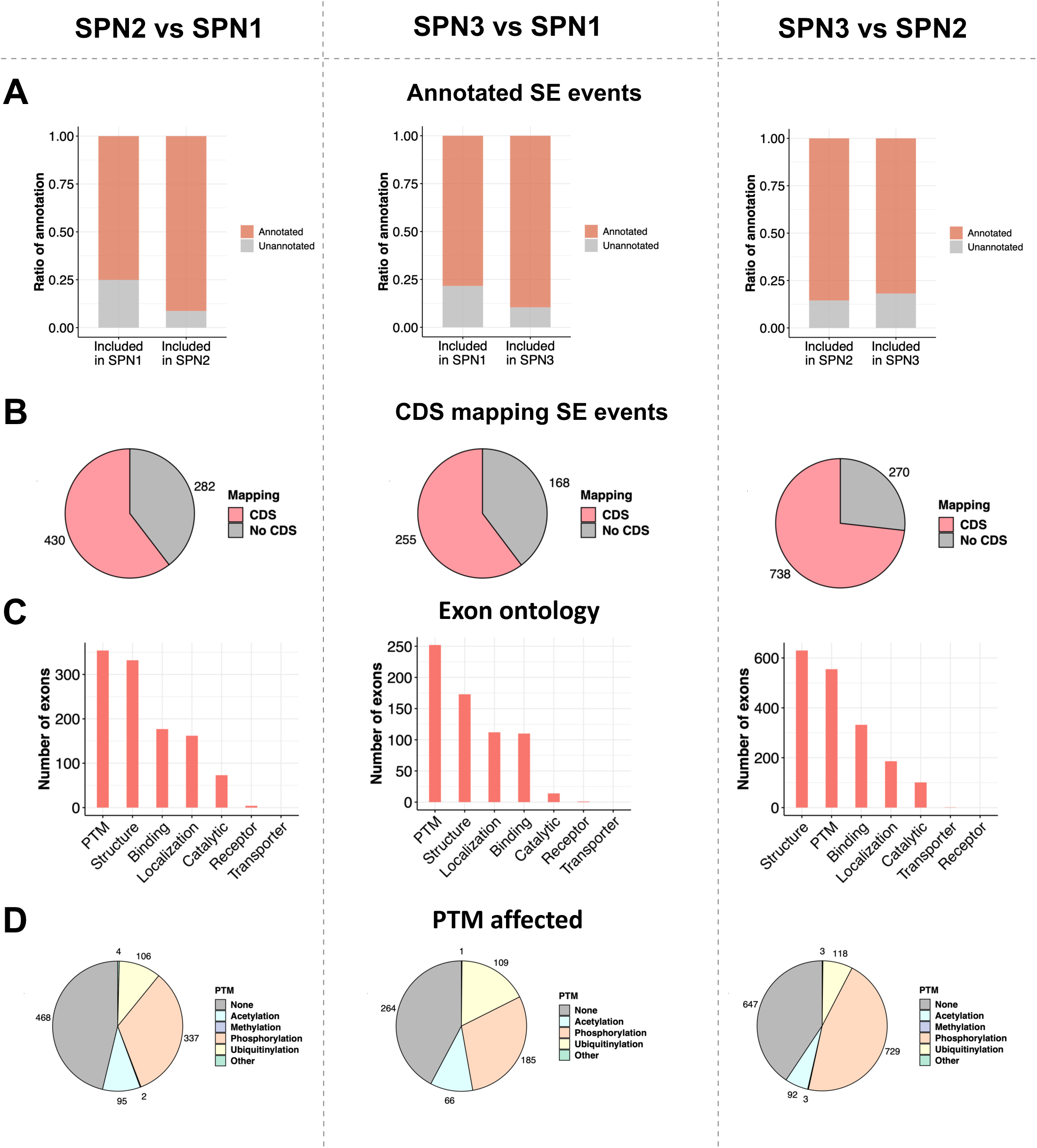
Analysis of differential skipped exons events when comparing SPN2 vs SPN1 (left), SPN3 vs SPN1 (center), and SPN3 vs SPN2 (right). **A** Bar plot illustrating the proportion of differential SE events that were previously annotated/unannotated. **B** Pie chart showing distribution of differential skipped exons in coding (CDS) and non-coding regions. **C** Bar plot showing the distribution of functional annotations for differential SE events. **D** Pie chart comparing the proportion of differential SE events affecting PTM sites.

**Supp Fig 5.**
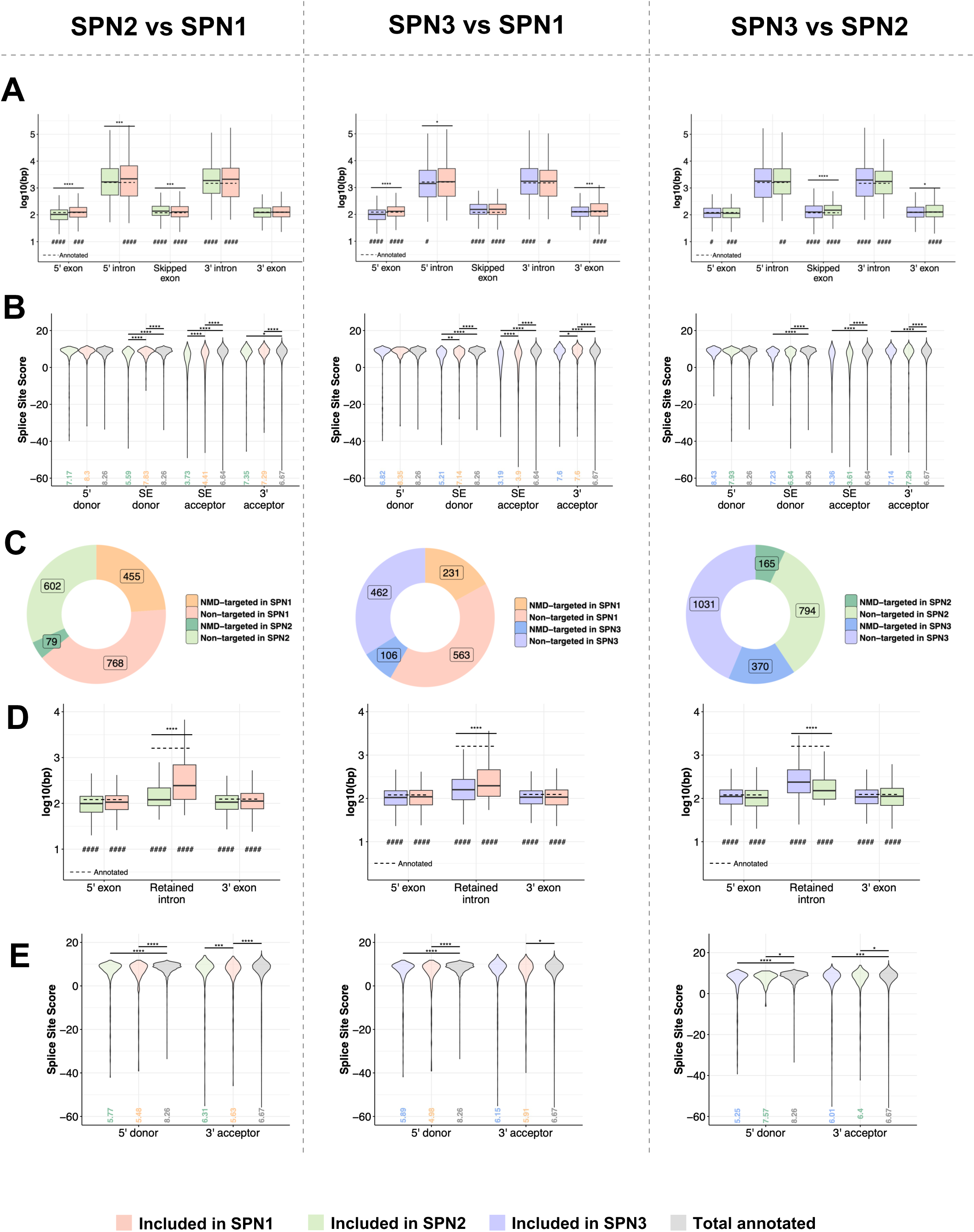
Analysis of exons and introns features in SE and RI events when comparing SPN2 vs SPN1 (left), SPN3 vs SPN1 (center), and SPN3 vs SPN2 (right). **A** Box plot showing the comparison of sizes of skipped exons and their upstream and downstream exons and introns for differential SE events. **B** Box plots depicting the distribution of splice site scores for donor and acceptor sites of differential SE events. **C** Pie chart illustrating the estimated proportion of differential SE events that lead to degradation by nonsense-mediated decay (NMD). **D** Box plot showing the comparison of sizes of retained introns and their upstream and downstream exons for differential RI events. **E** Box plots depicting the distribution of splice site scores for donor and acceptor sites of differential RI events.

**Supp Fig 6.**
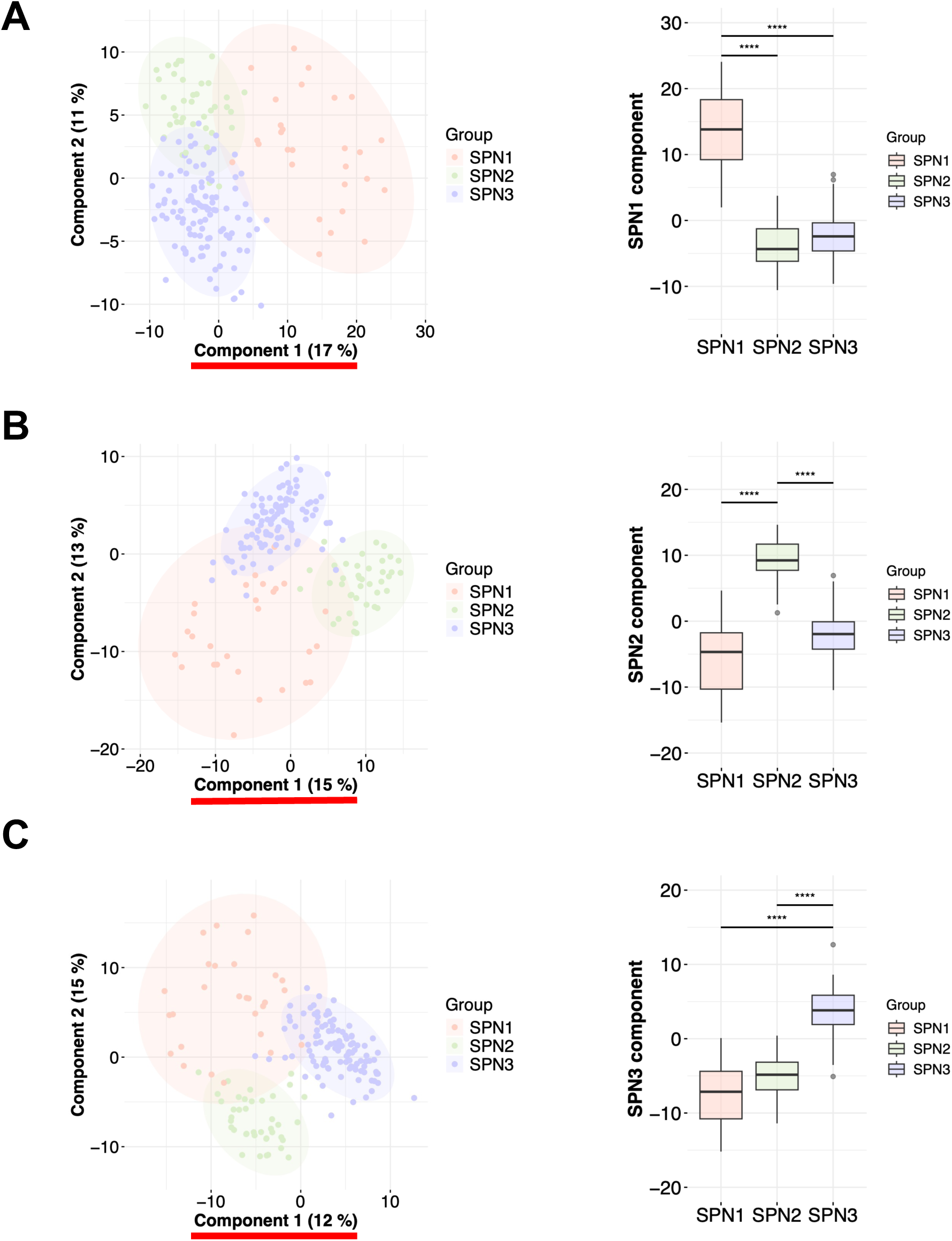
Partial Least Squares Discriminant Analysis (PLS-DA) clustering showing the separation of SPN1 (**A**), SPN2 (**B**) or SPN3 (**C**) against the rest, based on the expression profiles of splicing machinery genes. At the right, box plots showing the scores of the selected component for each group (underlined in red in the PLSDA plot). Asterisks indicate statistical significance of t test: **** *p* < 0.0001).

